# Dynamic control of Raf–ERK signaling modulates neuronal activity across biological scales

**DOI:** 10.64898/2026.01.07.698027

**Authors:** Huaxun Fan, Eunjoo Kang, Yuan Zhou, Collin Barnes, Nathan Gron, Hairuo Du, Catherine A. Christian-Hinman, Xinzhu Yu, Kai Zhang

**Affiliations:** Department of Biochemistry, University of Illinois at Urbana-Champaign, Urbana, Illinois 61801, USA; Neuroscience Program, University of Illinois at Urbana-Champaign, Urbana, Illinois 61801, USA; Department of Molecular and Integrative Physiology, University of Illinois at Urbana-Champaign, Urbana, Illinois 61801, USA; Present address: Center for Neuroimmunology and Glial Biology, The Brown Foundation Institute of Molecular Medicine for the Prevention of Human Diseases, The University of Texas Health Science Center at Houston, Houston, Texas 77030, USA; Science and Technology Center for Quantitative Cell Biology, University of Illinois at Urbana-Champaign, Urbana, Illinois 61801, USA

## Abstract

Neuronal activity robustly engages the extracellular signal-regulated kinase (ERK) signaling pathway through Ca^2+^-dependent mechanisms; however, whether ERK can acutely and causally modulates ongoing neuronal activity remains unsolved due to complex upstream regulation and diverse subcellular functions. Here, we directly address this question using an optogenetic ERK activator, opto-miniRaf, that enables selective, rapid, graded, and reversible control of ERK signaling. Combining this AAV-compatible system with calcium imaging and electrophysiology, we interrogate ERK functions across biological scales, from cultured neurons, acute brain slices, and the intact brain. Acute optogenetic activation of ERK enhances synchronized network burst activity in cultured rat cortical neurons and increases calcium activity of cortical pyramidal neurons in awake and moving mice following non-invasive light stimulation. Together, these results establish ERK signaling as an acute modulator of neuronal and network activity, positioning opto-miniRaf as a generalizable platform for precise spatiotemporal control of intracellular kinase signaling in complex biological systems.

## Introduction

Neuronal activity triggers a complex network of intracellular cascades that converge on the extracellular signal-regulated kinase (ERK) pathway. In neurons, synaptic input and membrane depolarization elevate intracellular Ca^2+^, activating small GTPases and kinases that drive ERK signaling (*1*). Through these routes, ERK has been extensively involved in activity-dependent processes across brain regions, from rapid biochemical responses to longer-term adaptations. As a result, ERK is implicated in not only sensory processing and motor control, but also in higher-level cognitive processes such as learning and memory (*2*). For example, ERK is activated in the dorsal hippocampus after behavioral training (*3*). By contrast, pharmacological inhibition of MEK, the upstream kinase regulator responsible for ERK activation, impairs hippocampal long-term potentiation(*4*), and ERK inhibition also leads to defective spatial learning in rodents (*5*).

Despite this well-established “neuronal activity to ERK” axis, it remains unclear whether ERK acutely modulates ongoing neuronal activity on the timescale of seconds to minutes. Dissecting this role has been challenging because of the promiscuity of ERK signaling. For example, the binding of neurotrophins to receptor tyrosine kinases (RTKs) often activates multiple parallel pathways, besides ERK, including the phosphatidylinositol-3 kinase/protein kinase B (PI3K/AKT) and phospholipase C-gamma (PLCγ) pathways (*6–8*). Similarly, GPCR activation of ERK leads to simultaneous activation of the cAMP-PKA axis. The other caveat is that pharmacological inhibitors, although widely used and thought to be specific, nonetheless elicit off-target effects that affect calcium entry to cells (*9*) or potassium channels (*10*). The dependence of ERK signaling outcomes on temporal kinetics (*11–14*) and spatial compartmentalization (*15*, *16*) adds additional challenges in dissecting ERK’s role in live neurons.

Optogenetics offers a powerful alternative, allowing for light-mediated control of cellular activity with high spatial and temporal resolution (*17*, *18*). Classical opsin-based tools, such as channelrhodopsins (*19*, *20*), enable precise manipulation of ion flow across the cell membrane, but do not directly engage intracellular signaling pathways. To address this challenge, several laboratories, including ours, have developed optogenetic tools to control intracellular signaling pathways. Among these, tools based on controlling the RTKs (*21–25*), chimeric GPCR (*26*, *27*), or GEF of Ras optoSOS (*28*), are capable of promoting ERK activation. However, these systems simultaneously activate multiple downstream signaling subcircuits. For example, activation of opsins promotes Ca^2+^ influx, which binds to calmodulin (CaM) and activates CaM kinases I, II, and IV. These kinases phosphorylate a wide array of substrates, including cytoplasmic proteins such as AKT, JNK, p38, and Rap1 (which activates ERK), as well as adenylate kinase (which triggers the cAMP–PKA pathway), and nuclear transcription factors, including CREB, SRF, and CBP. Optogenetically enabled GPCRs, such as optoXR (*26*) and opto-D1 (*27*), can activate the adenylate cyclase (AC)–cAMP–PKA cascade. PKA, in turn, phosphorylates a variety of downstream targets, including DARPP-32 (*29*) and ERK.

To delineate the direct signaling outcomes of ERK, we previously developed optoRaf, which bypasses upstream receptor activation by wiring a distinct route for membrane recruitment of Raf1 and selectively activating the Raf-MEK-ERK signaling cascade. OptoRaf successfully activates the Raf-MEK-ERK axis in cultured mammalian cell lines (*30*), *Xenopus laevis* embryos (*31*), and *Drosophila* larvae (*32*). Notably, optoRaf does not cross-activate other neurotrophic pathways, such as the PI3K-AKT or PLCγ pathways. However, optoRaf has not been used in mammalian brains due to challenges in effectively packaging transgenes in AAVs for in vivo delivery. To overcome this constraint, in this work, we engineered opto-miniRaf, an AAV compatible optogenetic system by systematically truncating Raf1 while preserving its ability to activate ERK. We demonstrated that opto-miniRaf enables selective and reversible ERK activation in cultured primary neurons, acute brain slices, and live mice. Acute ERK activation via opto-miniRaf enhances synchronized network activity in vitro and increases calcium activity in cortical pyramidal neurons in vivo. Together, opto-miniRaf provides a powerful platform for dissecting ERK-dependent regulation of neuronal activity across multiple physiological contexts.

## Results

### Screening for an AAV-compatible opto-miniRaf system

OptoRaf functions by light-mediated activation of Raf1 at the plasma membrane via blue-light-mediated interaction between photolyase homology region of cryptochrome 2 (CRY2PHR, abbreviated as CRY2 in this work) and (CIBN) at the plasma membrane (**Fig. 1A**) (*30–32*). Upon light-induced dimerization of CRY2 and CIBN, Raf1 is recruited to the plasma membrane. Whereas Raf1 membrane targeting can activate its activity (*33*), such activation is likely to be further enhanced by photo-oligomerization (tetramerization or formation of higher-order photobodies) of CRY2 (*34–37*). To expand its application and enable *in vivo*, neuron-specific expression, we aimed to integrate optoRaf with an AAV-based delivery system (*38*, *39*). A truncated calmodulin-dependent protein kinase II (CaMKII) promoter, CK0.4 (*40*), was selected to ensure neuron-specific expression of optoRaf. However, even after splitting the P2A-based single-plasmid system (*31*), the full length of core transgene CRY2-Raf1 (3.5 kb) and non-encoding elements, including inverted terminal repeats (ITRs, 290 bp), CK0.4 promoter (400 bp), WPRE (Woodchuck Hepatitis Virus Posttranscriptional Regulatory Element, 600 bp), and hGH polyA signal (480 bp), reaches 5.3 kb, exceeding the AAV packaging limit of 4.7 kb (*41*). These size constraints necessitated further optimization of CRY2-Raf1 for efficient AAV compatibility (**Fig. 1B**).

**Figure 1.**
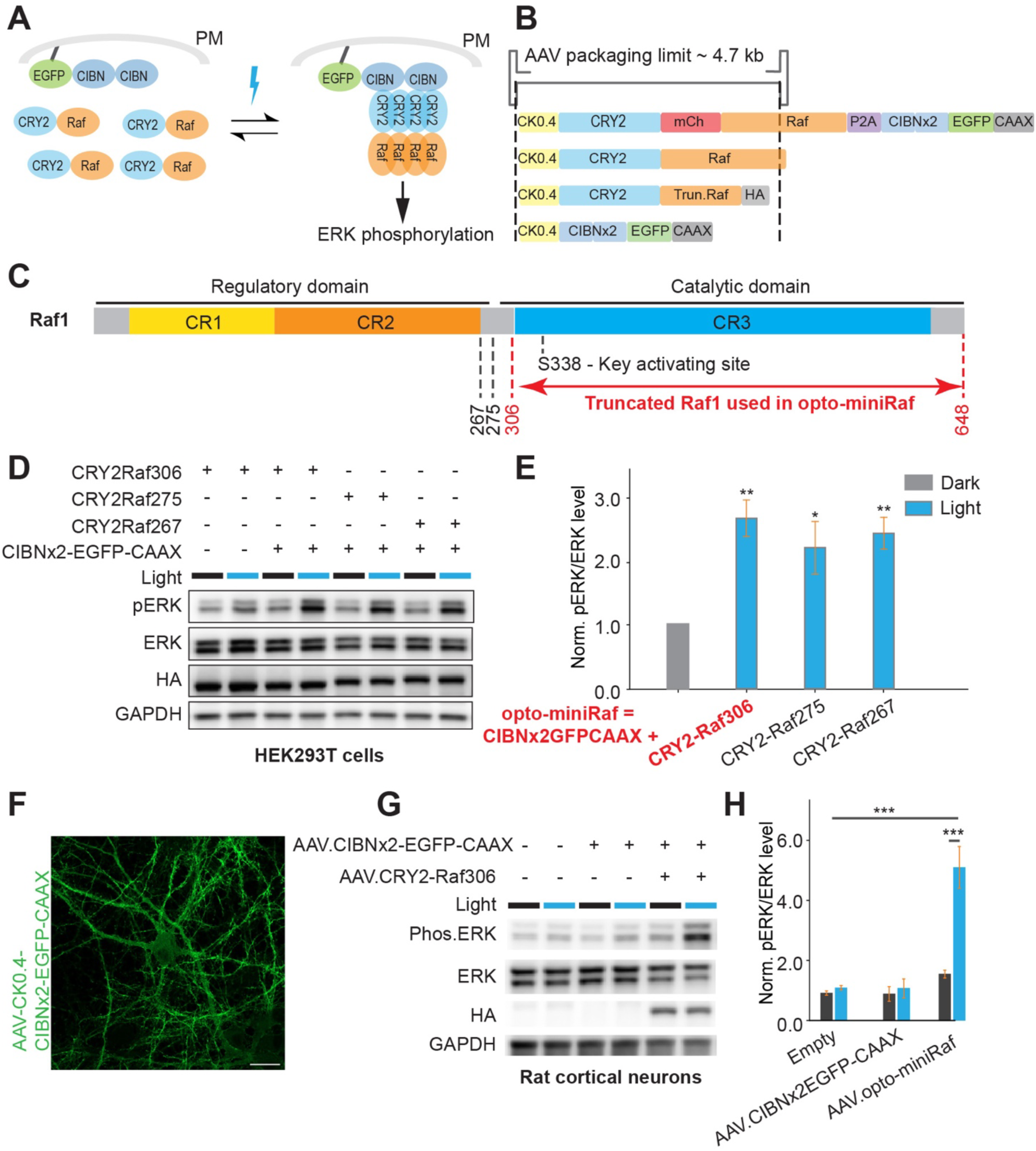
Screening for the opto-miniRaf for optogenetic activation of ERK signaling in primary rat cortical neurons. (A) Schematic of the opto-miniRaf system. Light-induced association of CRY2-Raf1 and CIBN×2-EGFP-CAAX recruits Raf1 to the plasma membrane (PM), where Raf1 likely undergoes clustering to activate the Raf/MEK/ERK signaling cascade. (B) Size of constructs with reference to the AAV packaging limit. (C) Domain structure of Raf1 and selection of truncation points, 267, 275, and 306. OptoRaf with truncated Raf1 fits into the AAV packaging limit. (D) Western blot analysis of HEK293T cells co-transfected with different CRY2-Raf1 truncations and CIBN×2-EGFP-CAAX following blue light stimulation (0.5 mW/cm^2^, 10 min). CRY2-Raf1(267-648), CRY2-Raf1(275-648), and CRY2-Raf1(306-648) were abbreviated as CRY2-Raf267, CRY2-Raf275, and CRY2-Raf306, respectively. (E) Quantification of normalized pERK/ERK levels from (D). pERK levels were normalized to total ERK. Samples maintained in the dark were defined as 1.0. (Unpaired one-tailed t-test was performed, n=3 biological replicates). (F) Representative confocal image of DIV13 primary rat cortical neurons transduced with AAV2/8.CK0.4-CRY2-Raf and AAV2/8.CK0.4-CIBN×2-EGFP-CAAX at DIV6. Scale bar: 20 μm. (G) Western blot analysis of primary rat cortical neurons transduced with AAV2/8.CK0.4-CRY2-Raf and AAV2/8.CK0.4-CIBN×2-EGFP-CAAX following blue light stimulation (0.5 mW/cm^2^, 10 min). Non-transduced neurons maintained in the dark served as a negative control. (H) Quantification of optoRaf activation in (G). PERK levels were normalized to total ERK. Non-transduced neurons maintained in the dark were defined as 1.0 (Unpaired one-tailed t-test was performed, n=3 biological replicates). *P<0.05, **P<0.01, ***P<0.001.

As the C-terminal Raf1 catalytic domain alone retains its kinase activity (*42*), we speculated that N-terminally truncated Raf1 should suffice for ERK activation in optoRaf. Truncation sites 267, 276, and 306 were selected based on a previous study (*43*) (**Fig. 1C**). To evaluate the performance of optoRaf with truncated Raf1, HEK293T cells were co-transfected with CIBN×2-EGFP-CAAX and different variants of CRY2-truncated Raf1. All three truncated variants induced ERK phosphorylation by more than 2-fold following 10 minutes of blue light stimulation at 500 µW/cm^2^, with the CRY2-Raf1 (306-648) showing the highest efficiency (**Fig. 1D-E**). Thus, we selected the CRY2-Raf1(306-648) as the final kinase component of opto-miniRaf. We also noticed that cells transfected with CRY2-Raf1(306-648) alone showed a slight increase in pERK upon blue light stimulation (**Fig. 1D**), likely due to CRY2-mediated Raf1 oligomerization in the cytoplasm. The final sizes of both transgenes, AAV-CIBN×2-EGFP-CaaX and CRY2-Raf1(306-640)-HA, are 1.8 kb and 2.6 kb, respectively, which allows for successful AAV packaging. When rat cortical neurons were transduced with AAV-optoRaf, which contains a 1:1 titer mixture of AAV-CK0.4-CIBN×2-EGFP-CaaX and AAV-CK0.4-CRY2-Raf1(306-640)-HA, on days in vitro (DIV) 6, steady expression of transgenes was observed three days post-transduction via Western blot analysis with primary antibody against the HA tag (**Fig. S1A**). Additionally, strong EGFP expression was detected in almost 100% of transduced neurons on DIV13 (**Fig. 1F**).

### Opto-miniRaf enables reversible ERK activation in vitro and ex vivo

To determine the performance of opto-miniRaf in neurons, we transduced rat cortical neurons with AAVs encoding opto-miniRaf on DIV6, followed by optogenetic stimulation with a 5-minute light stimulation (470 nm, 1 mW/cm^2^) on DIV13. This transient stimulation resulted in a 2.8-fold increase in the level of phosphorylated ERK (**Fig. 1G-H**). Extending the illumination duration from 5 to 10 minutes did not further enhance phosphorylation levels, indicating a rapid kinetics of ERK activation. A higher-power light stimulation at 5 mW/cm^2^ resulted in a slightly stronger increase in pERK levels (**Fig. S1B**).

To evaluate the optoRaf function in acute brain slices, stereotaxic injection(*38*) was performed to deliver AAV-opto-miniRaf into the mouse hippocampus (**Fig. 2A**). Significant EGFP fluorescence in the dentate gyrus region was observed at 3-4 weeks post-injection without the need for immunohistochemical amplification (**Fig. 2B**). To activate opto-miniRaf, an optical fiber coupled to a blue laser was positioned directly above acute brain slices. ERK activation proceeded in a dose-dependent manner when various stimulation durations from 1 to 5 minutes were applied (**Fig. 2C-D**). Control animals with no AAV injections showed no changes in pERK level upon blue light illumination (**Fig. 2C**). Furthermore, pERK levels declined after illumination ceased, returning to the baseline within 30 minutes (**Fig. 2E-F**). Together, these results demonstrate that opto-miniRaf enables acute ERK activation in primary rat cortical neuron cultures and acute mouse brain slices.

**Figure 2.**
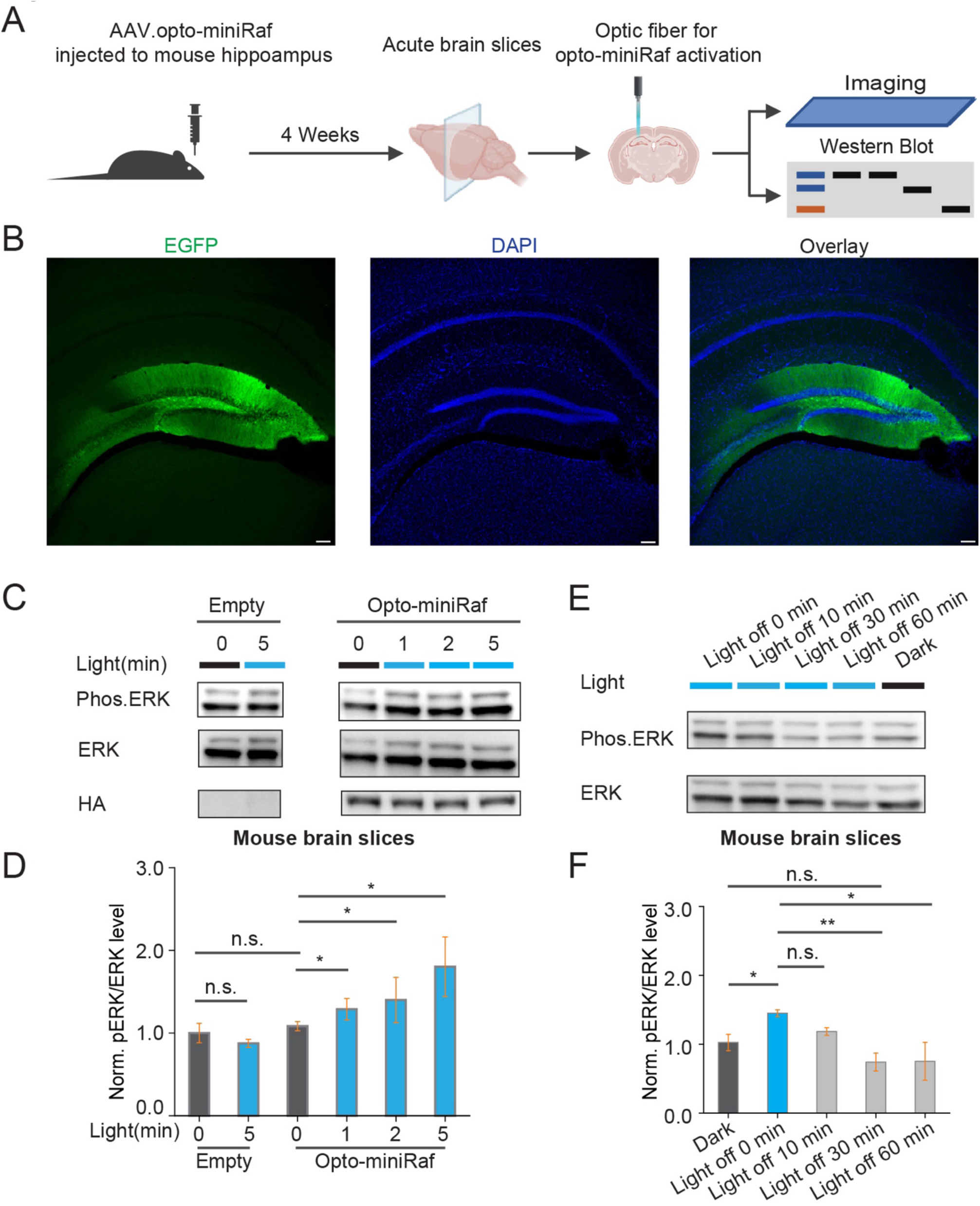
Opto-miniRaf reversibly activates ERK signaling ex vivo. (A) Schematic of the experimental setup. AAV2/8.CK (0.4)-CRY2-Raf and AAV2/8.CK(0.4)-CIBN×2-EGFP-CAAX were mixed at a 1:1 ratio and injected into the mouse hippocampus. Neurons transduced with both viruses are labeled as optoRaf in the figure. After 4 weeks of expression, acute brain slices were prepared, and an optical fiber was positioned above the hippocampus to deliver blue light stimulation. Slices were subsequently analyzed by imaging and Western blot. (B) Representative confocal images of AAV2/8.CK(0.4)-CIBN×2-EGFP-CAAX expression in fixed hippocampal sections. Scale bar: 100 µm. (C) Western blot analysis of brain slices following 1-, 2-, and 5-minute light stimulation (5 mW/cm^2^, 80 Hz, 90% duty cycle). (D) Quantification of pERK levels in (C). The pERK level was normalized to the total ERK level. Normalized pERK in non-injected animals maintained in the dark was defined as 1.0 (Unpaired one-tailed t-test was performed, n = n = 4 animals per condition, one brain slice per animal). (E) Western blot analysis of opto-miniRaf inactivation in brain slices 10, 30, and 60 minutes after cessation of light stimulation. (F) Quantification of pERK levels in (E). pERK levels were normalized to total ERK. Non-injected animals (no AAV) maintained in the dark were defined as 1(Unpaired one-tailed t-test was performed, n = 3 animals per condition, one brain slice per animal). *P<0.05, **P<0.01, ***P<0.001.

### Opto-miniRaf enhances spontaneous network burst activity in cultured cortical neurons

Next, we investigated whether optogenetic activation of ERK signaling modulates neuronal activity using fluorescence imaging of calcium dynamics in cultured rat cortical neurons. Cortical neurons were co-transduced at DIV 6 with AAVs encoding opto-miniRaf and jRCaMP1a, a genetically encoded calcium indicator that emits red fluorescence upon calcium binding (*44*). Robust neuronal expression of both constructs was observed by DIV 13 (**Fig. 3A**). Live-cell calcium imaging was then performed to record spontaneous network activity in these cultures. Prominent spontaneous calcium activity was observed under control conditions, including recurrent synchronized burst events involving a large fraction of recorded cells (**Fig. 3B; Video S1 & S2**), a characteristic feature of dissociated cortical neuronal networks (*45*). Accordingly, we focused on network-level synchronized activity as a robust readout of collective dynamics in vitro. Calcium traces were extracted from individual neurons using the Suite2p package (*46*), synchronized calcium burst activity was subsequently quantified as described in **Fig. S2**.

**Figure 3.**
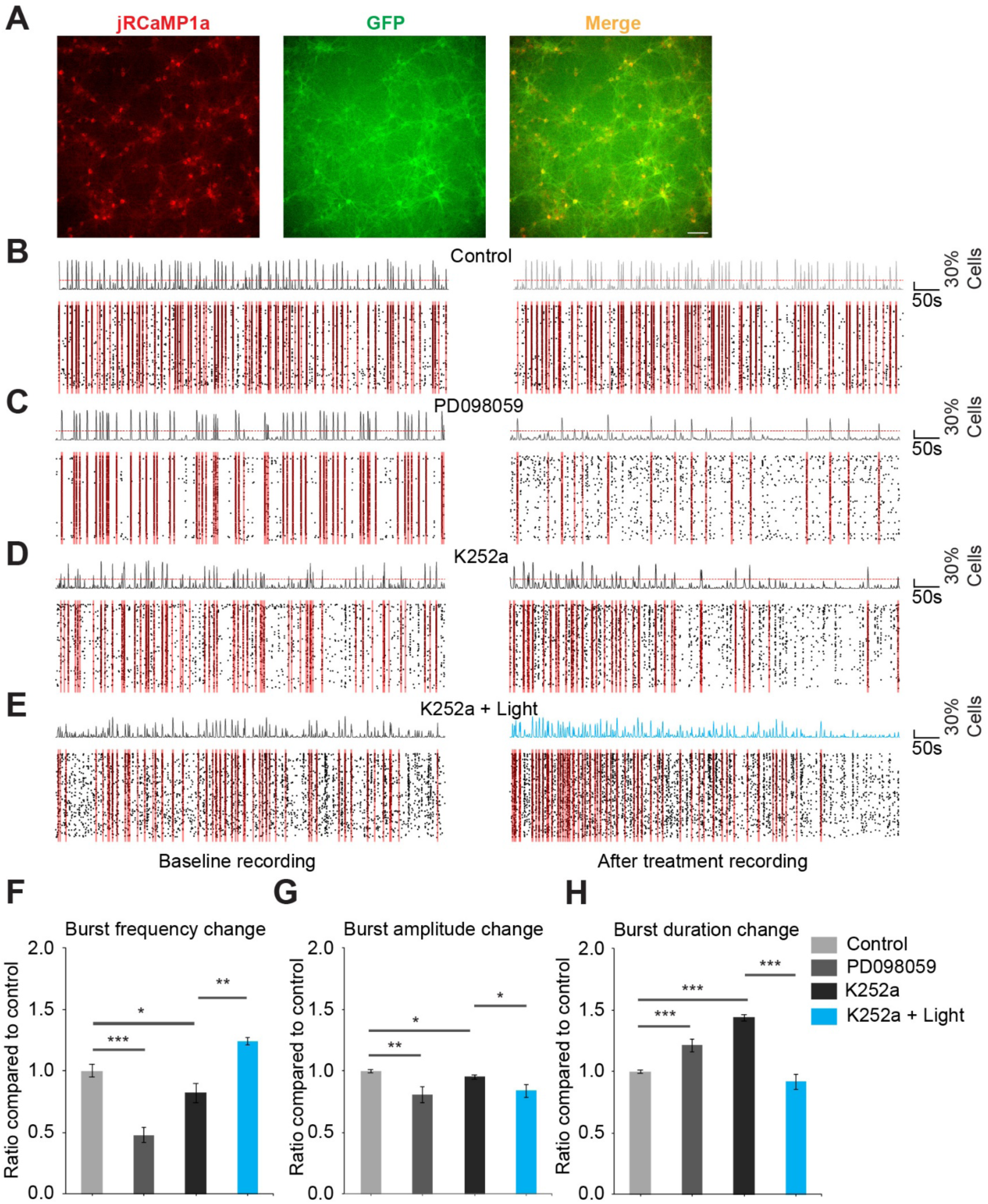
Opto-miniRaf modulate synchronized calcium burst activity in cultured cortical neurons. (A) Representative images of DIV13 cultured cortical neurons transduced with AAV opto-miniRaf and AAV jRCaMP1a at DIV6. Scale bar: 100 μm. (B) Representative raster plots of calcium events under control conditions. The left panel shows baseline activity, and the right panel shows activity after a 5-min rest period. Individual cell calcium events are shown as black ticks, with the population activity trace aligned above. The red dotted line indicates the synchronization threshold (30% of total recorded cells). Time periods exceeding this threshold are defined as synchronized burst events and highlighted with translucent red shading. *n* = 6 recording sets. (C) Same layout as in (B) for neurons treated with 2 μM PD098059. *n* = 8 recording sets. (D) Same layout as in (B) for neurons treated with 1 μM K252a. *n* = 4 recording sets. (E) Same layout as in (B) for neurons treated with 1 μM K252a, with blue-light stimulation (10 mW/cm^2^) applied immediately after completion of the baseline recording and maintained throughout the second recording period. *n* = 4 recording sets. (F–H) Quantification of synchronized burst properties across conditions. (F) Relative change in synchronized burst frequency, calculated as the ratio of post-treatment to baseline. (G) Relative change in synchronized burst amplitude. (H) Relative change in synchronized burst duration. Control groups are normalized to 1.0. Data are presented as mean ± SEM. Unpaired one-tailed t-test was performed. *P<0.05, **P<0.01, ***P<0.001.

To examine the contribution of ERK signaling to network synchronization, we first inhibited ERK activation using the MEK inhibitor PD098059, a well-characterized pharmacological blocker of MEK-dependent ERK activation (*47*) or K252a, a Trk receptor inhibitor (**Fig. 3C-E**) (*48*). Compared to control recordings, PD098059 (**Fig. 3C; Video S3& S4**) or K252a treatment (**Fig. 3D-E; Video S5-S8**) significantly altered synchronized burst activity, which is characterized by a marked reduction in burst frequency, a mild but significant reduction in burst amplitude, and a significant increase in burst duration (**Fig. 3F–H**). Notably, when opto-miniRaf was activated in the presence of K252a, all altered traits in burst activity were restored (**Fig. 3E–H**). As K252a blocks other neurotrophic downstream signaling pathways except for the Raf-MEK-ERK axis (activated by opto-miniRaf), these results indicate that ERK activation is solely responsible for the regulation of spontaneous synchronized network burst activity in cultured cortical neurons.

### Opto-miniRaf activation increases neuronal activity in the motor cortex of awake mice

To apply opto-miniRaf *in vivo*, we performed stereotaxic injections of AAVs encoding jRGECO1a and the two components of opto-miniRaf at a 1:1:1 titer ratio into the motor cortex, followed by transcranial window implantation for two-photon calcium imaging with two-photon microscopy (**Fig. 4A**). Immunohistochemistry for EGFP, jRGECO1a, and NeuN confirmed robust viral co-expression in the motor cortex neurons (**Fig. 4B**). Given that the transcranial window provides direct optical access to cortical tissue, a non-invasive external light source effectively activated opto-miniRaf in the motor cortex (**Fig. S3A**). To assess ERK activation, immunohistochemistry for pERK was performed (**Fig. 4C**). As expected, thirty minutes of light stimulation resulted in a 2.7-fold increase in pERK levels, while animals expressing opto-miniRaf but maintained in darkness or undergoing no AAV injection showed no change (**Fig. 4D**, **Fig. S3B-C**). Notably, ERK activation was observed in neurons located more than 1 mm beneath the cortical surface (**Fig. 4E**), highlighting the sensitivity of opto-miniRaf *in vivo*.

**Figure 4.**
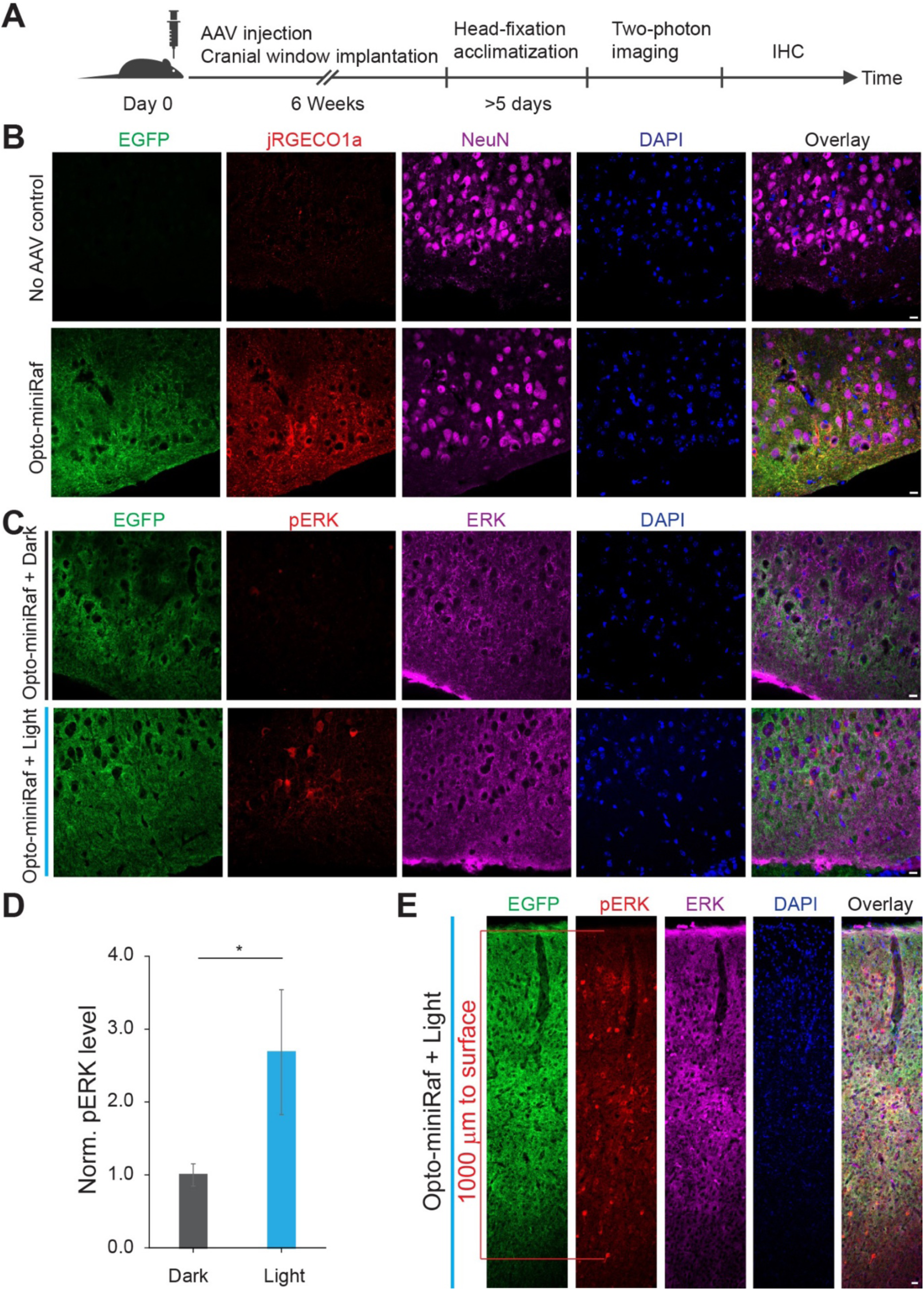
Opto-miniRaf activates ERK signaling in the mouse motor cortex. (A) Schematic of in vivo optogenetic and two-photon imaging experiments. AAV2/8.CK(0.4)-CRY2-Raf, AAV2/8.CK(0.4)-CIBN×2-EGFP-CAAX and AAV1/2.hSyn-NES-jRGECO1a-WPRE-SV40 were injected into the motor cortex of 8-10 week-old mice at a 1:1:1 ratio. After 6 weeks of expression, animals were habituated to head fixation before in vivo two-photon imaging. (B) Representative confocal images of EGFP, jRGECO1a, and NeuN immunostaining in brain slices from the AAV-injected hemisphere and the contralateral (uninjected) hemisphere. (C) Representative confocal images of EGFP, pERK, total ERK immunostaining and DAPI staining in brain slices from animals maintained in dark or stimulated with blue light (10 mW/cm^2^, 30 min). (D) Quantification of pERK levels in (C). pERK levels were normalized to total ERK, AAV-injected animals maintained in the dark were set as 1(Unpaired one-tail t-test was performed, n = 4 slices from 3 animals). (E) Representative extended-depth confocal images of EGFP, total ERK and pERK immunostaining in brain slices from animals stimulated with blue light (10 mW/cm^2^, 30 min). Scale bar: 20 μm. *P<0.05, **P<0.01, ***P<0.001.

To determine the functional consequences of opto-miniRaf activation, we next performed *in vivo* two-photon calcium imaging in awake, head-fixed animals. Two-photon imaging confirmed strong expression of both optoRaf components and jRGECO1a in the motor cortex (**Fig. S3D**). Fluorescence calcium traces were extracted from individual cortical neurons, and spike frequency and peak amplitude were quantified from raw ΔF/F_0_ signals (**Fig. 5A-B**). Notably, we monitored neuronal activity within the same neurons before and after light stimulation, which enabled the analysis of both ensemble-averaged and single-cell level responses to opto-miniRaf activation. We first confirmed that keeping animals in the dark for 30 min does not change the calcium spike frequency or amplitude, as shown by the spaghetti and histogram plot of the Dark/Baseline ratio of spike frequency and amplitude (**Fig. 5C**). In contrast, thirty minutes of light stimulation (10 mW/cm^2^) significantly increased both calcium spike frequency and amplitude, as shown by the spaghetti plot and the right shift of the histogram for Light/Baseline ratios (**Fig. 5D**). These results demonstrate that opto-miniRaf activation in the brain of head-fixed awake mice enhances neuronal activity.

**Figure 5.**
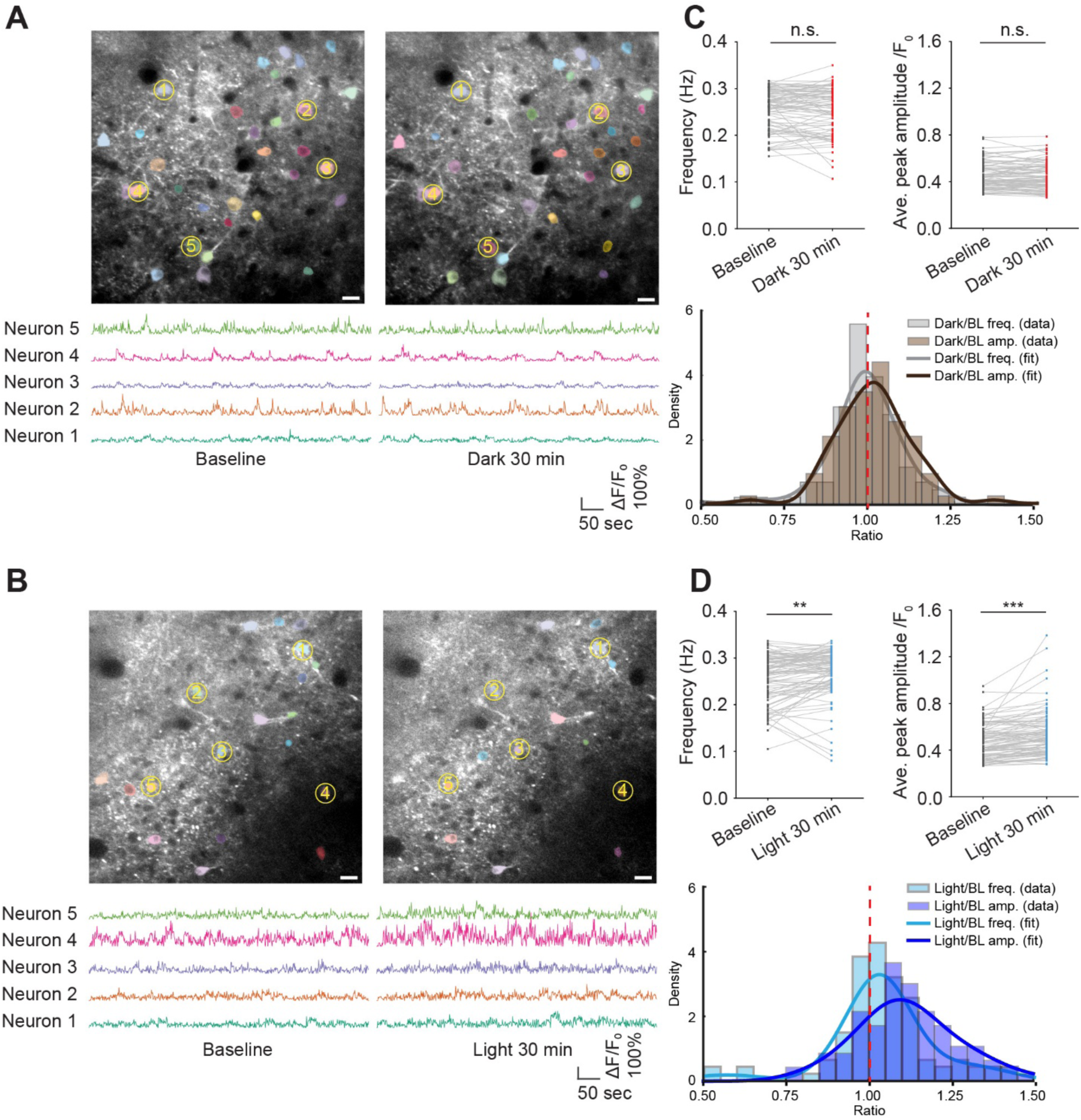
Opto-miniRaf activation in the mouse motor cortex increases neuronal activity. (A-B) Representative two-photon images of the same field of view (FOV) from baseline and after 30 minutes in the dark (A) or blue light (10 mW/cm^2^) (B), with ΔF/F_0_ traces for 5 example neurons (Yellow circled). (C-D) Quantified calcium activity frequency and average peak amplitude from (A) and (B). Peaks were defined as events exceeding 20% ΔF/F_0_ (Paired one-tailed t-test was performed, n=87 neurons from 6 animals, 7 image sets for dark group, n=95 neurons from 6 animals, 7 image sets for light group). Images are frame-averaged projections of 600 frames for visualization. Scale bar: 20 μm.

### OptoRaf activation dynamically modulates neuronal activity in freely moving animals

Taking advantage of the reversible ERK activation mechanism, we next evaluated the dynamic nature of ERK’s modulation on neuronal activity in freely moving animals. A paradigm for transiently activating opto-miniRaf in mice was developed (**Fig. 6A**). After a “baseline” recording for 10 minutes before any treatment, we applied a 1-minute two-photon optogenetic activation of optoRaf with a 940 nm pulsed laser, as it has been shown that CRY2 can be activated by two-photon excitation (*49*). A 10-minute “post-light” recording was performed immediately after optogenetic stimulation. The animals then rested in darkness for one hour, after which a final recording was taken (“Recovery”) (**Fig. 6A**). Notably, one minute of laser stimulation at 1 frame per second successfully increased the frequency and peak amplitude of calcium signaling. After one hour of recovery in the dark, calcium spike frequency and amplitude returned to baseline levels (**Fig. 6B-E**, **Fig. S4A-B**). In contrast, animals without opto-miniRaf AAV injection exhibited no change following 1-minute laser stimulation (**Fig. S5A-C)**). These results indicate that optoRaf can acutely and reversibly modulate neuronal activity in live animals.

**Figure 6.**
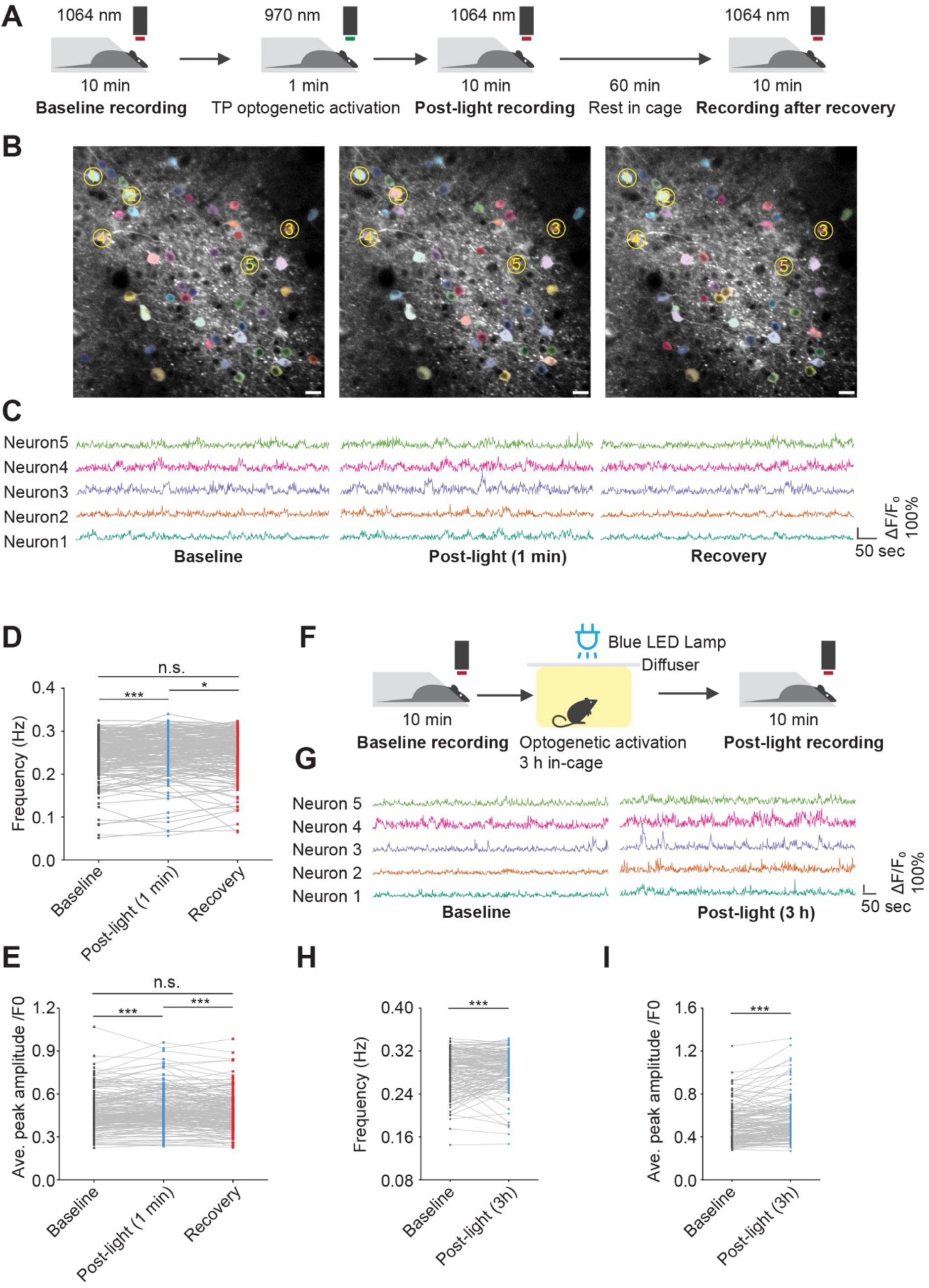
Opto-miniRaf acutely and reversibly modulates neuronal activity in the mouse motor cortex. (A) Schematic of experimental setup for 1 min laser activation. (B-C) Representative two-photon images (B) and traces (C) of the same field of view (FOV) from baseline, post-light(1min) and recovery, with ΔF/F_0_ traces for 5 example neurons (Yellow circled). Scale bar: 20 μm. (D-E) Quantified calcium spike frequency and average peak amplitude from (B-C) (Paired one-tailed t-test, n=196 neurons from 3 animals, 6 image sets). (F) Schematic of experimental setup for 3 h light stimulation in freely moving mice. (G) ΔF/F_0_ traces for 5 example neurons before and after 3 h blue light stimulation (10 mW/cm^2^). (H-I) Quantified calcium activity frequency and average peak amplitude from (G) (Paired one-tailed t-test, n=144 neurons from 7 animals, 7 image sets). Images are frame-averaged projections of 600 frames for visualization. Peaks were defined as events exceeding 20% ΔF/F_o_. *P<0.05, **P<0.01, ***P<0.001.

To investigate a more physiologically relevant light stimulation paradigm, freely moving animals were placed in a cage equipped with an overhead LED lamp, with a diffuser applied to ensure uniform light distribution (**Fig. 6F**). Three hours of illumination significantly increased both spike frequency and amplitude (**Fig. 6G-I**, **Fig. S6**). This finding highlights the potential of opto-miniRaf for studying the effects of ERK signaling activation in behaviorally relevant contexts.

## Discussion

In this study, we constructed opto-miniRaf, an AAV-compatible optogenetic system that enables light-inducible ERK activation *in vitro*, *ex vivo*, and *in vivo*. We validated this system in cultured primary rat cortical neurons and acute hippocampal slices, where brief blue light stimulation induced robust and reversible ERK phosphorylation. The light sensitivity of optoRaf enables effective ERK activation in the living mouse motor cortex using a non-invasive external light source, in contrast to the fiber optics commonly used in opsin-based systems. In cultured rat cortical neurons, optoRaf-induced ERK activation increased the frequency of synchronized neuronal activity. Furthermore, *in vivo* two-photon calcium imaging of the motor cortex revealed increased spontaneous calcium activity after light stimulation, consistent with enhanced neuronal activity.

Unlike other optogenetic systems based on opsins, RTKs, or GPCRs, opto-miniRaf bypasses ligand activation and selectively activates the ERK pathway with minimal crosstalk to other downstream pathways, as previously demonstrated (*32*). We are aware that N-terminally truncated Raf1 alone would be constitutively active. Indeed, as shown from the very first discovery of the retroviral oncogene, *v-raf*, a viral homolog of the cellular gene *Raf1*, truncation of Raf1’s CR1 and CR2 domains results in high basal activity independent of Ras activation (*42*). The N-terminus of Raf1 was then hypothesized to have two functions: binding to activated Ras, required for membrane recruitment, and inhibiting Raf1’s C-terminal kinase catalytic domain (*50*). In opto-miniRaf, i.e., CIBN×2-GFP-CaaX and CRY2-Raf(306-648), CRY2-CIBN interaction provides an alternative route of membrane recruitment under light stimulation. CRY2 could also exert an inhibitory effect on Raf1’s C-terminal catalytic domain, because only a slight increase in basal activity of CRY2-Raf(306-648) was observed in the dark (Figure 1H). We speculate that this inhibitory effect is released upon stimulation with blue light. Thus, CRY2 is likely to fulfill both functions of the N-terminal domains of Raf1 by using light as a new triggering mechanism in comparison to ligand-induced Raf1 activation.

We demonstrated that temporally controlled (down to 1 minute), mild-power (0.5-10 mW/cm^2^) blue light stimulation is sufficient to induce ERK phosphorylation. Upon removal of blue light illumination, ERK activity returns to the baseline within 30 min. In primary cortical neurons, opto-miniRaf activation enhances network-level synchronous calcium activity. In live mice, optoRaf stimulation effectively triggered acute ERK activation in motor cortex neurons, resulting in increased calcium signaling dynamics, as reflected in the elevated frequency and peak amplitude of calcium transients. Notably, the light sensitivity of opto-miniRaf enables the use of non-invasive, wide-field blue light (10 mW/cm^2^) through the implanted cranial window, allowing for fiber-less optogenetic stimulation of cortical areas in freely moving animals.

The increase in synchronized network activity observed upon opto-miniRaf activation raises important questions regarding the cellular mechanisms by which ERK signaling shapes population-level neuronal dynamics. Synchronized bursting in dissociated cortical networks is known to emerge from the interplay between intrinsic neuronal excitability and synaptic coupling within recurrent circuits (*51*, *52*). ERK signaling has been shown to regulate neuronal excitability through modulation of ion channel function, such as Kv4.2 potassium channels (*53*), as well as synaptic transmission and plasticity through trafficking of neurotransmitter receptors (*54*).

Given the system’s precise temporal control, opto-miniRaf is well-suited for dissecting the phase-specific roles of ERK signaling in cognitive processes, such as learning, memory consolidation, retrieval, and extinction. Previous studies have shown that neurotrophin delivery following behavior training enhances memory, and pharmacological inhibition of ERK activity completely abolishes this effect (*55*), highlighting ERK’s critical role in memory consolidation. However, pharmacological activation lacks spatiotemporal precision, and therefore, the mechanistic dissection of ERK at different memory phases is challenging. Previous studies using optoEphB2 in a fear conditioning paradigm found that light activation during the training phase enhanced long-term fear memory by 55% (*56*). Follow-up work can use opto-miniRaf to delineate the functional role of ERK signaling in fear memory formation among the multiple signaling subcircuits downstream of EphB2 signaling. Moreover, the light sensitivity of opto-miniRaf enables noninvasive light delivery in the superficial layer of the cortex, thereby minimizing behavioral disruption during in vivo experiments.

While the opto-miniRaf system offers high spatiotemporal modulation of ERK signaling, several limitations should be considered. First, our analyses were limited to spontaneous neuronal activity (*57*), which reflects baseline dynamics but does not reveal how ERK activation modulates stimulus-evoked responses and plasticity under defined input conditions. Future experiments incorporating evoked paradigms, such as sensory stimulation or optogenetic input, will be crucial in determining how ERK activity influences neuronal plasticity in more physiological contexts. Second, it would be advantageous to further simplify optoRaf into a single AAV system. Although it has been shown that CRY2-mediated Raf1(CRAF) homotypic association could lead to ERK activation (*58*, *59*), the resultant dynamic range is smaller than that of the current two-component optoRaf system under the same illumination conditions (Figure 1D), likely because CRY2 oligomerization requires a higher blue light power than CRY2-CIBN interaction, which would pose a challenge for in vivo applications.

By integrating pathway specificity, temporal precision, and in vivo applicability, opto-miniRaf provides a powerful platform for dissecting ERK signaling dynamics in the intact nervous system. This tool enables new opportunities to investigate the molecular basis of plasticity and behavior, which may inform future strategies for targeted neuromodulation in disease contexts.

## Materials and Methods

### Light stimulation for optogenetics in cultured neurons

For low-power light illumination on cultured neurons, a custom LED array was constructed by mounting 24 blue LEDs (Linrose Electronics, B4304H96) in a 6×4 configuration on a breadboard. Light intensity can be adjusted through a tunable power supply (Eventek, KPS3010D). The breadboard was housed in an aluminum box, and a light diffuser film was positioned above the LED array to ensure uniform illumination across the culture area. Light intensity at the plate of cell culture plate was measured using a power meter (Thorlabs, PM100D with S121C sensor). A stronger LED or a blue LED lamp (Utilitech, Par38) was used to achieve high-power (over 1 mW/cm^2^) light stimulation. The final light intensity was adjusted by changing the distance between the light source and the sample and tuning the power supply.

### Stereotaxic AAV injection in the mouse hippocampus

All animal experiments and procedures adhered to the guidelines set by the Institutional Animal Care and Use Committee of the University of Illinois at Urbana-Champaign. Adult C57BL/6J male and female mice (postnatal day 42 and older) were injected with AAV, performed following modifications of previously described procedures (*60*). The procedure was conducted under 2-3% oxygen-vaporized isoflurane anesthesia (Piramal Critical Care). On the day of injection, a 1:1 mixture of AAV-CK0.4-CIBN×2-GFP-CaaX and AAV-CK0.4-CRY2-RAF-HA was prepared (each diluted in sterile saline to a final 1×10^12^ titer). The final viral mixture was bilaterally injected (1 µL per side) into the dorsal hippocampus (coordinates: 2.0 mm posterior and 1.5 mm lateral to bregma; 1.4 mm ventral to the cortical surface) at a rate of 120 nL/min. After each injection, the needle was retained at the site for 3-5 min before removal. Subcutaneous carprofen (0.5 mg/mL, Division of Animal Resources, University of Illinois Urbana-Champaign) was administered at the beginning of surgery to manage pain. Additional intra- and postoperative care included the application of lubricant eye drops (Alcon Laboratories Inc.), 2.5 % lidocaine-2.5 % prilocaine cream (Alembic Pharmaceuticals ltd.), and Neosporin antibiotic ointment (Johnson and Johnson) to the wound after closing the incision with sutures (Ethicon).

### Acute brain slice preparation

Acute brain slices were prepared 3-4 weeks after AAV injection (or from non-injected controls) following established procedures (*60*, *61*). Briefly, after pentobarbital anesthesia (55 mg/kg i.p.), mice were euthanized by decapitation. The brain was rapidly excised, and coronal slices of 300-µm thickness were prepared using a Leica VT1200S vibratome. During slicing, the brain was immersed in an ice-cold, oxygenated (95% O_2_, 5% CO_2_), highly concentrated sucrose solution containing 234 mM sucrose, 11 mM glucose, 2.5 mM KCl, 0.5 mM CaCl_2_, 1.25 mM NaH_2_PO_4_, and 10 mM MgSO_4_. Immediately after slicing, each brain slice was hemisected along the midline, then incubated in oxygenated artificial cerebrospinal fluid (ACSF, containing 2.5 mM KCl, 10 mM glucose, 126 mM NaCl, 1.25 mM NaH_2_PO_4_, 1 mM MgSO_4_, 0.5 mM CaCl_2_, and 26 mM NaHCO_3_; Osmolarity ~ 298-300 mOsm) at 32°C for 1 hour, followed by transfer to room temperature (21-23°C) for at least 15 min before use.

### Light delivery to brain slices and tissue collection

For light stimulation of ex vivo brain tissue, individual slices were transferred to a chamber on the stage of an upright BX51WI microscope (Olympus America). The oxygenated ACSF in the chamber was heated to 30-32°C using an inline heater (Warner Instruments) and circulated at 2.5 mL/min. Blue light (473 nm) stimulation was delivered from a laser (Laserglow Technologies) via an optical fiber (FT200EMT, Thorlabs) placed directly above the surface of the slice, targeted to the dentate gyrus of the hippocampus. Blue light stimulation (20 Hz, 5 mW, 90% duty cycle)(*60*) was delivered to the slice for 5 min, and then the slices were held in the chamber for varying durations from 0 to 60 min after the end of the stimulation. Once post-light incubation was complete, slices were recovered from the chamber and briefly placed in a small dish of ACSF while the hippocampal tissue was manually dissected from the slice using a scalpel blade. The tissue was then placed in a microcentrifuge tube and rapidly frozen on dry ice. All samples were stored at −80°C until processing for western blot analysis.

### Calcium imaging in cultured primary cortical neurons

Primary cortical neurons were transduced at DIV5 with AAV.PHP.eB.Syn-NES-jRCaMP1a-WPRE-SV40, AAV.PHP.eB.CK(0.4)-CRY2-Raf1(306–648), and AAV.PHP.eB.CK(0.4)-CIBN×2-EGFP-CAAX. Control cultures received AAV.PHP.eB.Syn-NES-jRCaMP1a-WPRE-SV40 and AAV.PHP.eB.CK(0.4)-CIBN×2-EGFP-CAAX only. Live calcium imaging was performed at DIV13 using an inverted epifluorescence microscope (Leica DMI8) equipped with a 10× objective and an LED light source (SOLA SE II 365). Calcium signals were recorded using a Texas Red filter cube (Leica; excitation: 560/40 nm, dichroic: 595 nm, emission: 645/75 nm) at a frame rate of 4 frames per second. After live imaging, GFP expression was verified using a GFP filter cube (Leica; excitation: 472/30 nm, dichroic: 495 nm, emission: 520/35 nm) to avoid unintended optogenetic activation during validation.

### Surgical procedure for in vivo calcium imaging

All surgical procedures were performed under aseptic conditions using sterilized instruments. Adult mice were anesthetized with 4% isoflurane for induction and placed in a stereotaxic frame (David Kopf Instruments). Anesthesia was maintained at 1.25–2% isoflurane throughout the procedure, and body temperature was maintained at 37°C using a heating pad (RWD, Thermostat). Ophthalmic ointment (Terramycin) was applied to prevent corneal dehydration. Analgesic (Carprofen, 5 mg/kg) and anti-inflammatory (Dexamethasone, 0.2 mg/kg) agents were administered subcutaneously prior to surgery. The scalp was sterilized using alternating scrubs of betadine and ethanol, then incised to expose the skull above the motor cortex. A 3-mm circular craniotomy was made using a high-speed dental drill (Henry Schein Inc.), centered at ~+0.8 mm anterior and ±1.5 mm lateral to bregma. Artificial cerebrospinal fluid (ACSF; 124 mM NaCl, 4.5 mM KCl, 1.2 mM NaH_2_PO_4_, 26 mM NaHCO_3_, 10 mM D-glucose, 1 mM MgCl_2_, 2 mM CaCl_2_) was applied continuously to assist in skull thinning and to protect the exposed brain tissue during drilling.

Following the removal of the skull flap, the cortical surface was kept submerged in ACSF. A pre-pulled glass pipette (1B100-4; World Precision Instruments) filled with AAV was lowered into the motor cortex (coordinates: AP +0.8 mm, ML ±1.5 mm, DV –0.5 mm from the pial surface). AAVs were injected at a rate of 100 nL/min using a microinjection pump (UMP3; World Precision Instruments). The pipette was left in place for 10 minutes after the injection to minimize backflow. After viral delivery, a sterilized 3-mm glass coverslip was placed over the craniotomy and secured with tissue adhesive (VetBond). A lightweight titanium head-bar was then affixed to the skull, and the remaining exposed areas were sealed with dental cement. Postoperative care included recovery on a heating pad until full ambulation was achieved, with access to food and water. Carprofen and dexamethasone were administered subcutaneously once daily through postoperative day 4 (POD 4). Amoxicillin was provided in the drinking water for one week following surgery to prevent infection.

### In vivo two-photon imaging

Following a six-week recovery period after cranial window implantation, mice were gradually acclimated to head fixation over five consecutive days by incrementally increasing fixation duration. Once habituated, awake imaging was performed using an Ultima 2P two-photon microscope (Prairie Technologies/Bruker) equipped with a 20× 1.0 NA water-immersion objective (Olympus). Excitation wavelengths of 970 nm and 1064 nm (Coherent Axon laser) were used to visualize EGFP and jRGECO1a, respectively. Vascular and cellular landmarks were used to consistently relocate the same imaging region across sessions. Imaging was conducted at 1 Hz, with a field of view of approximately 320 μm × 320 μm.

### Immunohistochemistry of fixed mouse brain tissues

Mice were deeply anesthetized with isoflurane and transcardially perfused with 1× phosphate-buffered saline (PBS), followed by 10% formalin. Brains were post-fixed in 10% formalin overnight at 4°C and then transferred to 30% sucrose in PBS for cryoprotection. After full saturation, the brains were sectioned at a thickness of 40 μm using a cryostat (CM3050 S; Leica). Free-floating sections were incubated overnight at 4°C with the following primary antibodies diluted in PBS containing 0.3% Triton X-100 and 5% normal goat serum. Primary antibodies used include Chicken anti-GFP (1:1000 Abcam Cat. ab13970, RRID: AB_300798), Rabbit anti-RFP (1:1000, Rockland Cat. 600-401-379, RRID: AB_2209751), Mouse anti-NeuN (1:1000, Millipore Cat. MAB377, RRID: AB_2298772), Rabbit anti-pERK (1:400, Cell Signaling Technology Cat. 4370, RRID: AB_2315112), Mouse anti-ERK (1:200, Cell Signaling Technology Cat. 4696, RRID: AB_390780). The following day, sections were washed three times with PBS and incubated for 2 hours at room temperature with fluorophore-conjugated secondary antibodies (1:1000). Secondary antibodies used include Alexa Fluor 488 goat anti-chicken (Molecular Probes Cat. A-11039, RRID: AB_142924), Alexa Fluor 594 goat anti-rabbit (Thermo Fisher Scientific Cat. A-11037, RRID: AB_2534095) and Alexa Fluor 647 goat anti-mouse (Thermo Fisher Scientific Cat. A-21235, RRID: AB_2535804). After washing, sections were mounted onto glass slides with antifade mountant with DAPI (Invitrogen, P36962). Confocal fluorescent images were acquired using a Zeiss LSM900 confocal laser scanning microscope (Carl Zeiss, Germany) through a 40× oil-immersion objective. Excitation wavelengths were selected as 405 nm for DAPI, 488 nm for GFP, 561 nm for RFP, 647 nm for CY5, and emission was collected using spectral detectors with standard filter settings. Images were acquired under identical acquisition parameters across both experimental and control groups to facilitate quantitative comparison.

## QUANTIFICATION AND STATISTICAL ANALYSIS

Quantification of Western blot bands and analysis of fluorescence images were conducted using ImageJ. For calcium imaging analysis, fluorescence traces from individual neurons were extracted from two-photon recordings using the Suite2p package (*62*). Raw fluorescence traces corresponding to each region of interest (ROI) were converted to ΔF/F_0_ traces using a custom Python script. The baseline fluorescence (F_0_) was defined as the minimum fluorescence intensity within a sliding 20-second window for each time point. Calcium events were identified as local maxima with a minimum inter-peak distance of 1 frame and a threshold of 20% ΔF/F_0_. Event frequency and average peak amplitude were calculated using the same custom analysis pipeline. For paired comparisons before and after stimulation, only neurons that could be reliably tracked and matched across both conditions were included. Details of statistical tests, exact n values, and what each n represents are provided in the corresponding figure legends. Statistical significance was defined as follows: P < 0.05 (*), P < 0.01 (**), and P < 0.001 (***). Data analysis was performed using Microsoft Excel and Origin Pro.

## Supporting information

VideoS1_Control_Baseline

VideoS2_Control_PostTreatment

VideoS3_PD098059_Baseline

VideoS4_PD098059_PostTreatment

VideoS5_K252a_Baseline

VideoS6_K252a_PostTreatment

VideoS7_K252a_Light_Baseline

VideoS8_K252a_Light_PostTreatment

## Funding

National Institute of General Medical Sciences R01GM132438 (KZ)

National Institute of Mental Health R01MH124827 (KZ)

National Science Foundation Science and Technology Center for Quantitative Cell Biology 2243257 (KZ)

Cancer Center at Illinois 9572 (KZ)

National Institute of Mental Health R01 MH132556 (XY)

National Institute of Health DP2 NS136871D (XY)

Whitehall Foundation research grant 2021-08-025 (XY)

Brain and Behavior Research Foundation NARSAD Young Investigator grant 30748 (XY)

National Institute of Neurological Disorders and Stroke R01 NS105825 (CAC-H)

## Author contributions

Conceptualization, HF, KZ, XY and CAC-H

Methodology, HF, EK, CAC-H. and YZ

Investigation, HF, EK, CAC-H, YZ, CB, NG and HD

Writing—original draft, HF, KZ, XY, CAC-H, EK and YZ

Writing—review & editing, KZ, XY and CAC-H

Funding acquisition, KZ, XY and CAC-H

Supervision, KZ, XY and CAC-H

## Data and materials availability

All data are available in the main text or the supplementary materials. Plasmids generated in this study are available upon request from the lead contact, provided a completed Materials Transfer Agreement is submitted. All original code is available upon request from the lead contact.

