## Supplementary Materials for "Dynamic control of Raf–ERK signaling modulates neuronal activity across biological scales"

**This PDF file includes:**

Supplementary Text
Supplementary Figures S1–S6

**Other Supplementary Materials for this manuscript include the following:**

Video S1-S8, Calcium imaging for primary cortical neurons, related to Figure 3. Videos were included in separate files.

### Supplementary Text

#### Plasmids construction

Truncated versions of Raf1 (residues 267–648, 275–648, and 306–648) were generated by PCR amplification from the previously published construct pEGFP-N1.CRY2-mCh-Raf1(S338A)-P2A-CIBN×2-EGFP-CAAX. Truncated CRY2-Raf1 fragments were subcloned into the pCS2 vector under a cytomegalovirus (CMV) promoter using In-Fusion Snap Assembly (Takara Bio, 638948) for expression in mammalian cells. For AAV-compatible constructs, CRY2-Raf1(306–648) and CIBN×2-EGFP-CAAX were independently cloned into the AAV-CK(0.4)GW[40] backbone (Addgene, 27226) using In-Fusion Snap Assembly. PCR amplifications were performed using the SimpliAmp Thermal Cycler (Applied Biosystems, A24811). PCR products were purified using the GeneJET Gel Extraction Kit (Thermo Scientific, K0692). All final constructs were sequence-verified by ACGT, Inc. or Plasmidsaurus.

#### Mammalian cell culture and transfection

HEK293T cells were grown in Dulbecco's modified Eagle Medium (Corning, 10-013-CV) supplemented with 10% fetal bovine serum and penicillin and streptomycin (Corning, 30-002-CI) in a humidified 5% $CO_2$ incubator at 37 °C. For transfection, cells were prepared with 40-60% confluency in various plates. The total volume of the medium and transfection mixture was scaled based on the plate area. For a 12-well plate, cells were incubated with 1 mL of growth medium and transfected with 1 µg of plasmid in 100 µL of transfection mix, which contained Turbofect (ThermoFisher, R0532) and a DNA mixture with a ratio of 1 µg DNA: 3 µL Turbofect. The mix was incubated for 20 min at room temperature and dispensed dropwise to the cell plate. The plate was gently rocked to aid even diffusion of the mix. After three hours of incubation, the medium was replenished with the complete culture medium. Downstream assays were performed at the desired time.

#### Western blot

Following light stimulation, neurons were washed once with 1 mL cold Dulbecco's PBS (DPBS) and lysed in 100 µL of cold RIPA buffer (Millipore, #20-188) supplemented with Halt™ Protease and Phosphatase Inhibitor Cocktail (Thermo Scientific, #78442). Lysates were centrifuged at 17,000 × *g* for 10 minutes at 4°C, and the supernatant was collected. Protein concentrations were determined using the Bradford assay (Thermo Scientific, #23238) and equalized across samples. Normalized lysates were mixed with LDS sample buffer (Invitrogen, NP0007), boiled for 5 minutes at 95°C, and loaded onto 10% polyacrylamide SDS-PAGE gels(Bio-Rad, 4561033). Electrophoresis was performed at room temperature, and proteins were transferred to PVDF membranes(Bio-Rad, 1620177) overnight at 30 V and 4°C. Membranes were blocked in TBST with 5% BSA for 1 hour at room temperature and then probed with primary and secondary antibodies according to the manufacturer's guidelines. The following primary antibodies were used: ERK(Cell Signaling Technology Cat. 4695, RRID: AB_390779), pERK(Cell Signaling Technology Cat. 4370, RRID: AB_2315112), HA-Tag(Cell Signaling Technology Cat. 3724, RRID: AB_1549585), GAPDH(Cell Signaling Technology Cat. 2118, RRID: AB_561053). An HPR-linked goat anti-rabbit secondary antibody(Cell Signaling Technology Cat. 7074, RRID: AB_2099233) was used. Membranes were incubated with ECL substrate and imaged using Azure 400 imager. Signal intensity analysis was performed with Fiji[56].

**AAV production**

For in vivo application, customized AAVs encoding CK0.4-CYR2-Raf and CK0.4-CIBN×2-EGFP-CAAX were produced at Boston Children's Hospital's viral core with serotypes 2/8. AAV2/1-Syn-NES-jRGECO1a-WPRE-SV40 was directly purchased from Addgene (100854-AAV1). For primary neuron application, AAVs were produced in-house, where HEK293T cells were transfected with the AAV helper plasmid pAdDeltaF6 (Addgene, 112867), the AAV cap plasmid pUCmini-iCAP-PHP.eB[57] (Addgene, 103005) and the AAV vector plasmids carrying optogenetic systems pAAV-CK(0.4)-CYR2-Raf, pAAV-CK(0.4)-CIBN×2-EGFP-CAAX, or calcium sensor pAAV-Syn-NES-jRCaMP1a-WPRE-SV40[44] (Addgene, 100848). After transfection, the cells were maintained in serum-free DMEM for 24 hours to recover before being switched to complete growth medium. After 3 days of viral production, both cells and culture medium were collected. The mixture was centrifuged at 800 rpm for 5 min, and the pellet was resuspended and homogenized via pipetting. The homogenate and supernatant were combined and filtered through a 0.22 μm membrane filter (Millipore, SLGS033). The filtrate was then incubated with DNase I (New England Biolabs, M0303) for 1 hour at 37°C to degrade free nucleic acids. Finally, the virus-containing solution was concentrated using a 100 kDa molecular weight cutoff concentrator (Millipore, UFC9100) to further increase the virus titer. Viral genome copies (GC) were quantified by qPCR using primers specific to the AAV ITR region, following DNase I treatment and proteinase K(New England Biolabs, P8107S) digestion to release encapsulated genomes.

**Primary neuron culture and AAV transduction**

Primary cortical neurons were prepared from embryonic day 19(E19) rat embryos. Timed-pregnant rats were euthanized in a $CO_2$ chamber and decapitated to obtain embryos. Brains were extracted, and cortical tissue was dissected after the removal of the meninges. Cortices were incubated in 3 mg/mL *Aspergillus melleus* protease (Sigma, P4032) at 37°C for 10 min, then dissociated into single cells via gentle trituration with flame-polished Pasteur pipettes. Cell counts were determined using a hemocytometer (Millipore, Z359629). Cells were then plated onto poly-D-lysine (Sigma, P1149)-coated 12-well plates or 12 mm coverslips in 24-well plates at densities scaled to the surface area. Cells were seeded in plating media composed of MEM (ATCC, 30-2003) supplemented with 10% FBS, 0.45% (w/v) D-glucose, 25 μM L-glutamine, 1 mM sodium pyruvate, and penicillin/streptomycin. 3 hours after initial seeding, the plating medium was replaced with maintenance medium consisting of Neurobasal medium supplemented with B27plus (Gibco, A3653401), 0.5 mM L-glutamate and penicillin/streptomycin. Cultures were maintained at 37°C in a humidified incubator with 5% $CO_2$. On DIV5, half of the medium was switched to fresh maintenance medium containing 2 μM cytosine β-D-arabinofuranoside (Ara-C; Sigma, C6645) to inhibit glial proliferation. For prolonged culture, the medium was subsequently refreshed twice per week by replacing half the volume with fresh maintenance medium. AAV transduction was performed at DIV 5 during the medium switch. Viruses were diluted in fresh maintenance medium and added to the culture at an estimated multiplicity of infection (MOI) of 5,000 to 10,000. Downstream analysis was performed at the desired time.

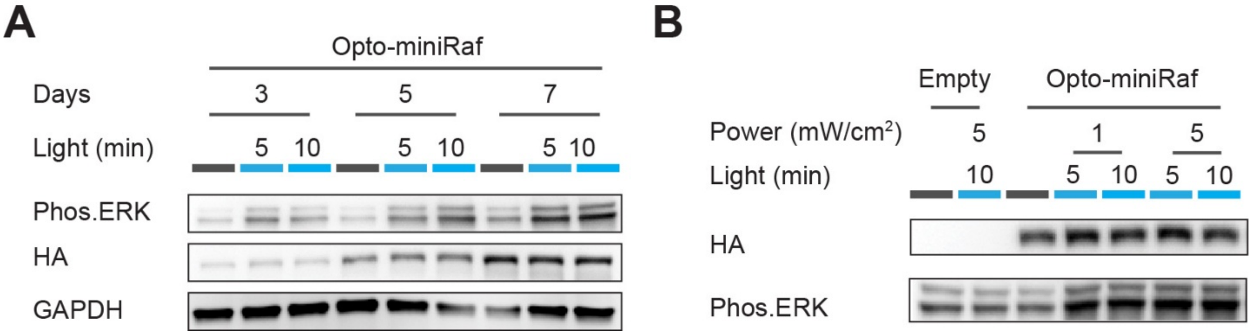


**FigureS1. Opto-miniRaf induces light-dependent ERK activation in primary cortical neurons. (A)** Western blot analysis of primary rat cortical neurons transduced with AAV.CK0.4-optoRaf following blue light (0.5 mW/cm$^2$) stimulation on various days post-transduction. Neurons maintained in the dark served as negative controls. **(B)** Western blot analysis of primary rat cortical neurons transduced with AAV.CK0.4-opto-miniRaf following stronger blue light (1 mW/cm$^2$, 5 mW/cm$^2$) stimulation. Activation was assessed after 5 min and 10 min of stimulation. Non-transduced neurons exposed to 5 mW/cm$^2$ blue light for 5 min served as a negative control.

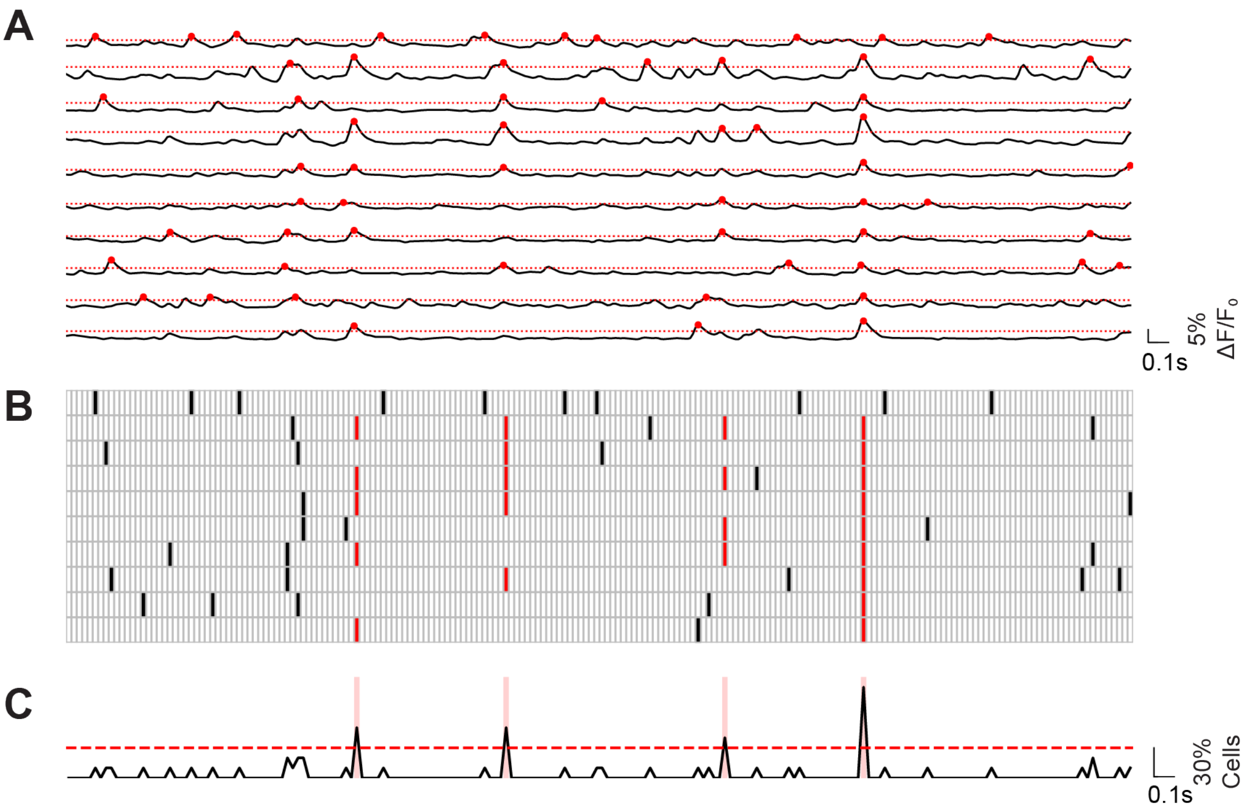


**FigureS2. Average calcium imaging trajectories for cultured rat cortical neurons. (A)** Representative ΔF/F traces from 10 individual neurons during the first 100 s of recording. Cell-specific activity thresholds are indicated by red dotted lines and detected calcium events are marked with red dots. **(B)** Binary raster plot generated from the traces shown in (A) after temporal binning at 500 ms resolution. Individual cell activity is shown in black, and synchronized burst events across the population are highlighted in red. **(C)** Population-based activity trace calculated as the fraction of active cells per time bin. The red dotted line indicates the synchronization threshold (30% of total cells). Collective activities beyond this threshold are defined as synchronized burst events, highlighted with translucent red shading.

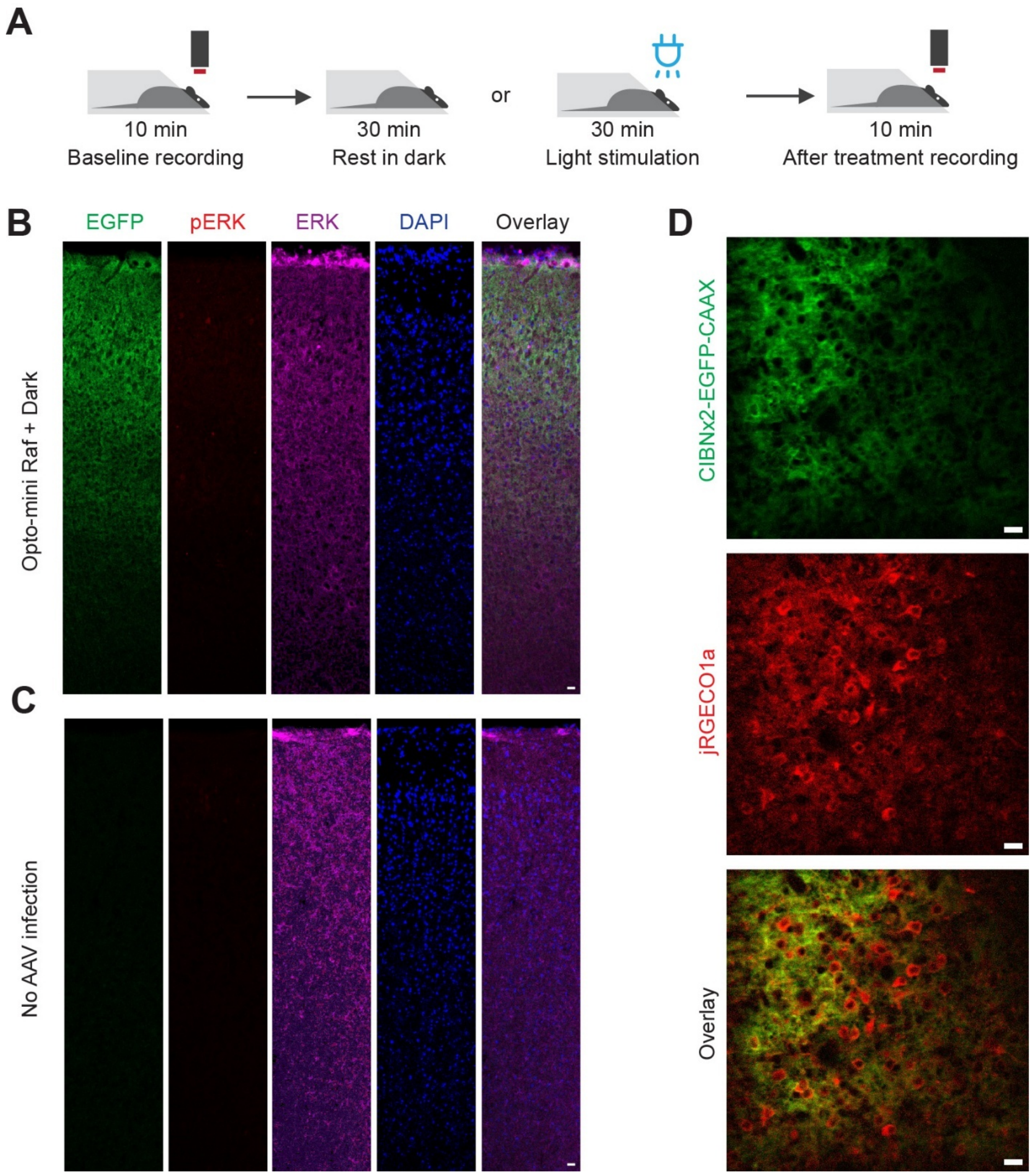


**Figure S3 In vivo opto-miniRaf activation and two-photon calcium imaging in the motor cortex.** (**A**) Schematic of in vivo experiments. Following a 10-minute baseline recording, animals remained head-fixed and were either kept in the dark or exposed to blue light (10 mW/cm²) for 30 minutes. A second 10-minute imaging session was performed immediately after light treatment. **(B)** Representative confocal images of DAPI, EGFP, pERK, and ERK immunostaining in brain sections from optoRaf-injected hemisphere with no light stimulation. **(C)** Images from the contralateral (non-injected) motor cortex. **(D)** Representative two-photon images showing expression of CIBN×2-EGFP-CAAX and jRGECO1a in the motor cortex. Images are frame-averaged projections of 60 frames for visualization. Scale bar: 20 μm.

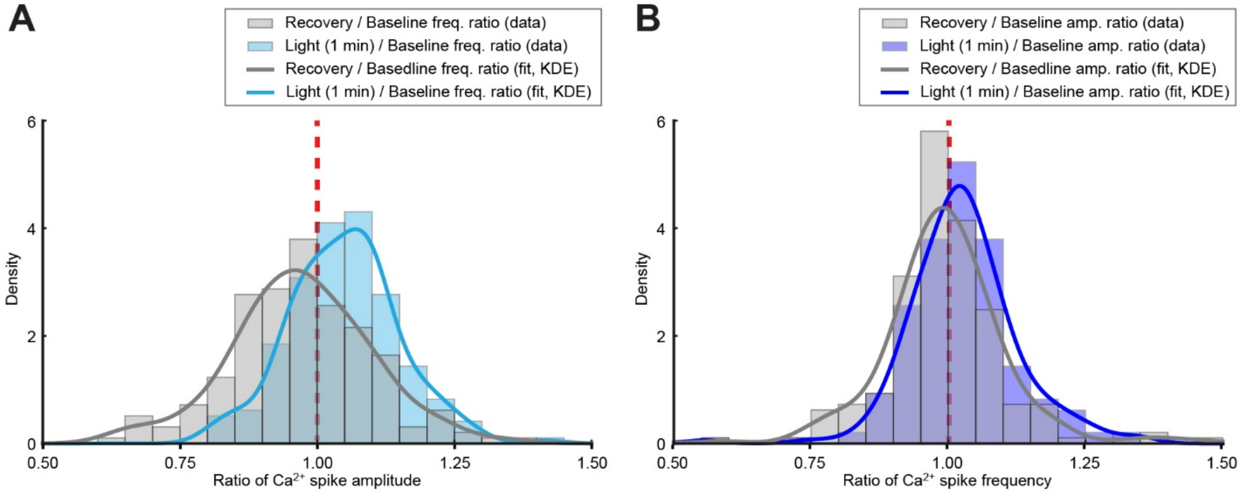


**Figure S4. Quantification of calcium spike frequency and amplitude. (A)** Histogram and KDE fitting of the ratio of calcium spike amplitude for Light/Baseline (cyan) and Recovery/Baseline (gray). **(B)** Histogram and KDE fitting of the ratio of calcium spike frequency for Light/Baseline (blue) and Recovery/Baseline (gray). Light illumination was performed for 1 min on a two-photon excitation confocal microscope.

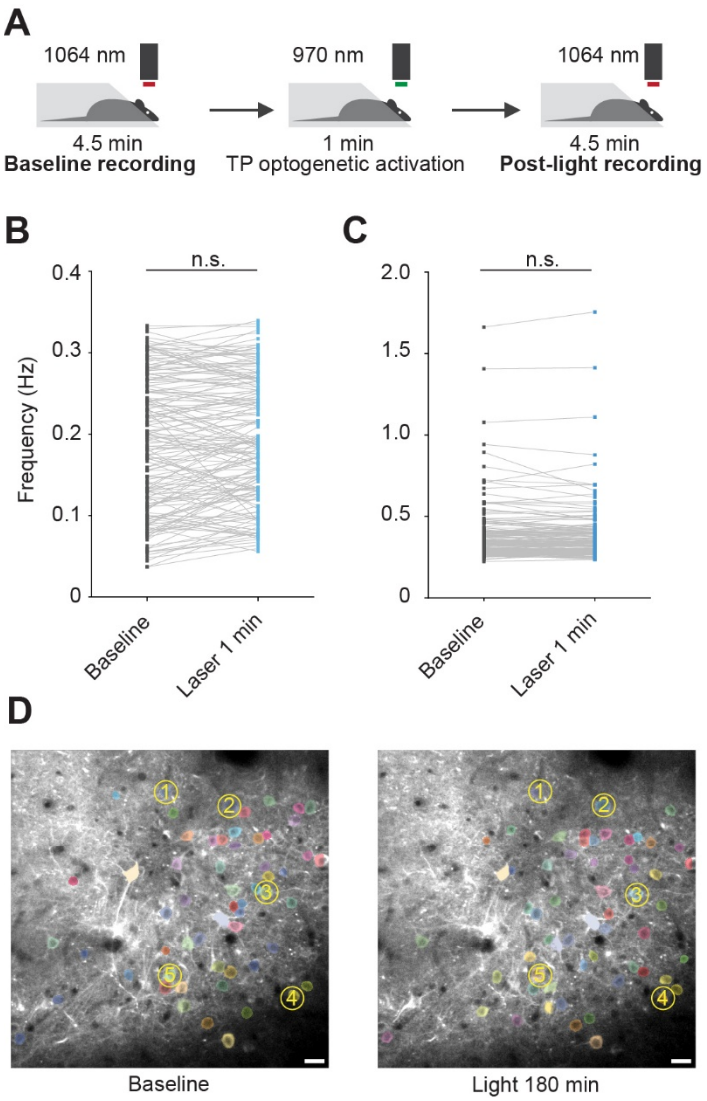


**Figure S5. Blue light illumination alone does not change neuronal excitability in the mouse motor cortex. (A)** Schematic of light stimulation two-photon imaging process. Following a 4.5-minute baseline recording, animals were exposed to 970 nm laser stimulation for 1 min at 1 frame per second(fps). A second 4.5-minute imaging session was performed immediately after light treatment. Animals injected with AAVs encoding the calcium sensor alone were used in this experiment (no optoRaf expression). **(B)** Quantified calcium activity frequency and average peak amplitude from (A). Peak were defined as events exceeding 20% $\Delta F/F_0$ (Paired two-tail *t*-test was performed, *n*=168 neurons from 3 animals 5 image sets). **(C)** Representative two-photon images of the same FOV from baseline, immediately after 1-minute laser stimulation, and after a 60-minute recovery period. **(D)** Representative two-photon images of the same FOV from baseline and after 3 hours of ambient blue light stimulation in the cage. Scale bar = 20 µm. *$P$<0.05, **$P$<0.01, ***$P$<0.001.

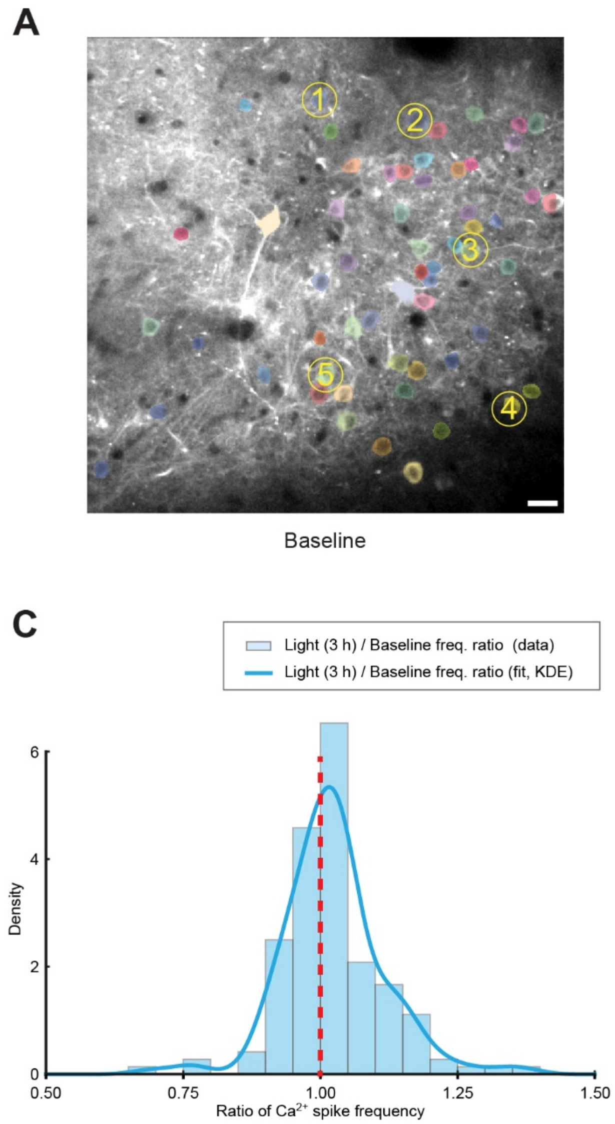


**Figure S6. Quantification of calcium spike frequency and amplitude after 3 h of opto-miniRaf activation in freely moving animals. (A-B)** Representative images of identical field of views from live mice with motor cortex transduced with AAV.opto-miniRaf and AAV.jRGECO1a before (A) and after 3 h of light illumination (B). **(C)** Histogram and KDE fitting of the ratio of calcium spike frequency of Light (3 h illumination in moving animals)/Baseline. **(D)** Histogram and KDE fitting of the ratio of calcium spike amplitude of Light (3 h illumination in moving animals)/Baseline.

**Supplementary Movie S1**
Live-cell calcium imaging for primary cortical neurons transduced with opto-miniRaf before dark incubation. Scale bar: 100 μm.

**Supplementary Movie S2**
Live-cell calcium imaging for primary cortical neurons transduced with opto-miniRaf after 5 min dark incubation. Scale bar: 100 μm.

**Supplementary Movie S3.**
Live-cell calcium imaging for primary cortical neurons transduced with opto-miniRaf before treatment with 2 μM PD098059. Scale bar: 100 μm.

**Supplementary Movie S4.**
Live-cell calcium imaging for primary cortical neurons transduced with opto-miniRaf after treatment with 2 μM PD098059 for 5 min. Scale bar: 100 μm.

**Supplementary Movie S5.**
Live-cell calcium imaging for primary cortical neurons transduced with opto-miniRaf before treatment with 1 μM K252a. Scale bar: 100 μm.

**Supplementary Movie S6.**
Live-cell calcium imaging for primary cortical neurons transduced with opto-miniRaf after treatment with 1 μM K252a for 5 min. Scale bar: 100 μm.

**Supplementary Movie S7.**
Live-cell calcium imaging for primary cortical neurons transduced with opto-miniRaf before treatment with 1 μM K252a and light stimulation. Scale bar: 100 μm.

**Supplementary Movie S8.**
Live-cell calcium imaging for primary cortical neurons transduced with opto-miniRaf after treatment with 1 μM K252a and blue light stimulation for 5 min (10 $mW/cm^2$ at the sample plane). Blue light was maintained throughout the recording. Scale bar: 100 μm.
